# Neural inflammation alters synaptic plasticity probed by 10 Hz repetitive magnetic stimulation

**DOI:** 10.1101/2020.10.16.336065

**Authors:** Maximilian Lenz, Amelie Eichler, Pia Kruse, Andreas Strehl, Silvia Rodriguez-Rozada, Itamar Goren, Nir Yogev, Stefan Frank, Ari Waisman, Thomas Deller, Steffen Jung, Nicola Maggio, Andreas Vlachos

## Abstract

Systemic inflammation is associated with alterations in complex brain functions such as learning and memory. However, diagnostic approaches to functionally assess and quantify inflammation-associated alterations in synaptic plasticity are not well-established. In previous work, we demonstrated that bacterial lipopolysaccharide (LPS)-induced systemic inflammation alters the ability of hippocampal neurons to express synaptic plasticity, i.e., the long-term potentiation (LTP) of excitatory neurotransmission. Here, we tested whether synaptic plasticity induced by repetitive magnetic stimulation (rMS), a non-invasive brain stimulation technique used in clinical practice, is affected by LPS-induced inflammation. Specifically, we explored brain tissue cultures to learn more about the direct effects of LPS on neural tissue, and we tested for the plasticity-restoring effects of the anti-inflammatory cytokine interleukin 10 (IL10). As shown previously, 10 Hz repetitive magnetic stimulation (rMS) of organotypic entorhino-hippocampal tissue cultures induced a robust increase in excitatory neurotransmission onto CA1 pyramidal neurons. Furthermore, LPS-treated tissue cultures did not express rMS-induced synaptic plasticity. Live-cell microscopy in tissue cultures prepared from a novel transgenic reporter mouse line [*C57BL6-Tg(TNFa-eGFP)*] confirms that *ex vivo* LPS administration triggers microglial tumor necrosis factor alpha (TNFα) expression, which is ameliorated in the presence of IL10. Consistent with this observation, IL10 hampers the LPS-induced increase in TNFα, IL6, IL1β, and IFNγ and restores the ability of neurons to express rMS-induced synaptic plasticity in the presence of LPS. These findings establish organotypic tissue cultures as a suitable model for studying inflammation-induced alterations in synaptic plasticity, thus providing a biological basis for the diagnostic use of transcranial magnetic stimulation in the context of brain inflammation.

## INTRODUCTION

Alterations in cognitive function and behavior are often observed in the context of systemic inflammation and/or infection of the central nervous system [1; 2; 3]. Several immune mediators that affect the ability of neurons to express plasticity have been identified [4; 5]. This is of considerable relevance in the context of neurological and psychiatric diseases associated with increased levels of pro-inflammatory cytokines in the brain [6; 7; 8; 9]. However, diagnostic approaches to functionally assess and quantify inflammation-associated alterations in neuronal plasticity and to monitor the effects of (plasticity-restoring) therapeutic interventions as yet remain underdeveloped.

Transcranial magnetic stimulation (TMS) is a promising clinical tool in this context. Based on the principle of electromagnetic induction, TMS allows for the non-invasive stimulation of cortical brain regions through the intact skin and skull [10]. The effects of single TMS pulses over the primary motor cortex can be quantified by recording motor evoked potentials in the target muscle [11]. Consecutive trains of TMS pulses (repetitive TMS, rTMS) over the motor cortex promote enduring changes of cortical excitability, i.e., neural plasticity [12]. For example, high-frequency rTMS (≥ 5 Hz) typically increases cortico-spinal excitability, measured as an increase in the amplitude of motor-evoked potentials [13; 14; 15]. Pharmacological approaches and analogies to basic research findings provide evidence to suggest that high-frequency rTMS asserts its effects via ‘long-term potentiation (LTP)-like’ plasticity [16; 17]. Previous studies employing animal models of rTMS (both in vitro and in vivo) have revealed that rTMS is capable of inducing long-lasting changes in glutamatergic neurotransmission that are consistent with the LTP of excitatory neurotransmission [18; 19; 20; 21; 22]. Since rTMS induces plasticity phenomena, in the present study, we hypothesize that this form of plasticity will be affected by neural inflammation.

In a previous study we reported that bacterial lipopolysaccharide (LPS)-induced systemic inflammation affects the ability of neurons to express LTP at the Schaffer collateral-CA1 synapses in the hippocampus [23]. While it is suspected that LPS can act directly on neuronal tissue to induce inflammation and alterations in synaptic plasticity, direct evidence for this remains scarce. Furthermore, the major source of the pro-inflammatory cytokine tumor necrosis factor alpha (TNFα), which has been linked to synaptic plasticity modulation [24; 25; 26; 27] is not well-characterized, while recent data suggest a distinct role of microglia derived TNFα in the regulation of LTP [28]. Therefore, in the present study, we employed *in vitro* LPS-exposure of entorhino-hippocampal tissue cultures prepared from a novel TNFα-reporter mouse strain that expresses the enhanced green fluorescent protein (eGFP) under the control of the *Tnfa* promoter [*C57BL/6-Tg(TNFa-eGFP)*] to monitor LPS-induced neural inflammation. Following our previous work on the cellular and molecular effects of repetitive magnetic stimulation (rMS) [19; 21; 29] and based on our recent findings of TNFα-mediated changes in synaptic plasticity [30], we tested whether excitatory synaptic plasticity induced by 10 Hz rMS is affected by *in vitro* LPS-induced inflammation, and whether the anti-inflammatory cytokine interleukin 10 (IL10) restores rMS-induced synaptic plasticity.

## MATERIALS AND METHODS

### Ethics statement

Mice were maintained in a 12 h light/dark cycle with food and water available *ad libitum*. Every effort was made to minimize distress and pain of animals. All experimental procedures were performed according to the national animal welfare legislation and approved by the animal welfare committee (V54-19c20/15-F143/37), the local animal welfare officers (Universities of Frankfurt, Mainz and Freiburg) and/or the institutional animal care and use committee (Sheba Medical Center).

### *C57BL/6-Tg(TNFa-eGFP)* reporter mouse line

To create a genetic construct that allows for the visualization of TNFα expression through simultaneous eGFP expression, the bacterial artificial chromosome (BAC) RP23-115O3 (GenBank: CR974444.18, BACPAC resources center at the children’s hospital Oakland research institute [CHORI]) was used as follows: Rabbit β-globin intron (Ac. No.: J00659; nucleotides: 907-1550) was amplified by the use of polymerase chain reaction (PCR) and cloned into pBluescript KS^+^ (Stratagene) via ClaI-EcoRI sites. The eGFP construct/sequence fused to the SV40 early mRNA polyadenylation (PA) signal was generated by applying standard PCR technique, using pEGFP-N1 (clontech) as a template and appropriate primers to delete the NotI site and cloned downstream to the Rabbit β-globin intron via EcoRI-SalI sites (all primers are given in the supplementary information, Table S1). The murine *Tnfa* promoter sequence including the first 167 nucleotides of exon 1 (Ac. No.: Y00467.1; nucleotides: 3120-4526) was inserted upstream to the rabbit β-globin intron via NotI-HindIII sites. A PGK-EM-neo cassette flanked by two full FRT sites was amplified using appropriate primers and the YTT vector B (Ac. No.: AY028413.1; nucleotides: 410-2310) as a template using standard PCR protocol and cloned as a SalI-NheI fragment downstream to the SV40 PA sequence. Finally, the DNA coding sequence immediately after the *Tnfa*’s PA signal (BAC RP23-115O3; nucleotides: 135366-134412) was obtained by PCR amplification and cloned downstream to the neo cassette via NheI-KpnI sites. The BstBI-KpnI fragments containing the coding sequences of the *Tnfa* promoter, the eGFP construct, the neo cassette and the *Tnfa* downstream region were used to replace the *Tnfa* coding sequence (nucleotides: 137835 to 135366) at BAC RP23-115O3 by the use of Red ET homologous recombination methods [31]. Positive BAC clones were analyzed by PCR using the upstream (U1-2a) and downstream (D1-2a) primers. BACs that were cloned correctly were further evaluated by southern blotting using the 5’-and the 3’-probes. Finally, the 5’- and the 3’-recombination junctions were sequenced to verify the correct integration sites of the reporter construct into the *Tnfa* locus. The recombinant eGFP reporter BAC was linearized using the restriction enzyme BsiWI (Thermo scientific) and purified using a CL4b sepharose column (Pharmacia). Eluted DNA was diluted to 1 µg/ml in an injection buffer (10 mM Tris, 0.1 mM EDTA, 100 mM NaCl, pH 7.5 with HCl) and injected into pronuclei oocytes (C57BL/J6-DBA hybrid). Transgenicity of the founders was verified by standard PCR methods using primer pairs corresponding to the Rabbit β-globin intron and eGFP sequence. The eGFP reporter efficiency was tested *in vitro* and *in vivo* and three founder strains were selected for additional experiments. Finally, to avoid disturbance of the neo gene, transgenic mice were bred with *FLP* mice (Jackson laboratory) and the neo-deleted offspring was backcrossed to *C57BL/6J* mice. Transgenic progeny was born at a normal Mendelian ratio and showed no signs of developmental abnormality or inflammatory insult.

### Preparation of tissue cultures

Organotypic entorhino-hippocampal tissue cultures were prepared at postnatal day 3–5 from *C57BL/6J* (Charles River) and heterozygous *C57BL/6-Tg(TNFa-eGFP)* reporter mice of either sex as previously described [32]. Culturing medium contained 50% (v/v) MEM, 25% (v/v) basal medium eagle, 25% (v/v) heat-inactivated normal horse serum (NHS), 25 mM HEPES buffer solution, 0.65% (w/v) glucose, 0.15% (w/v) bicarbonate, 0.1 mg/ml streptomycin, 100 U/ml penicillin and 2 mM glutamax. The pH was adjusted to 7.30 and tissue cultures were incubated for at least 18 days at 35°C in a humidified atmosphere with 5% CO_2_. Culturing medium was replaced by fresh medium three times per week.

### Pharmacology

Tissue cultures (≥ 18 days in vitro) were treated with lipopolysaccharide (1 µg/ml; LPS from Escherichia coli O111:B4, Sigma #L4391) for 3 days (c.f., [33]), which induced a robust increase in *Tnfa*-mRNA levels in our preparations (Figure 3). In some experiments, interleukin 10 (10 ng/ml; mouse recombinant, R&D Systems #417-ML) was added to the culturing medium. Age- and time-matched control cultures were treated with vehicle-only in the same way.

### Live-cell imaging

Live-cell imaging of heterozygous *C57BL/6-Tg(TNFa-eGFP)* cultures was performed at a Zeiss LSM800 microscope equipped with a 10x water-immersion objective (NA 0.3; Carl Zeiss). Filter membranes with 3 to 6 cultures were placed in a 35 mm Petri Dish containing pre-oxygenated imaging solution consisting of 50% (v/v) MEM, 25% (v/v) basal medium eagle, 50 mM HEPES buffer solution (25% v/v), 0.65% (w/v) glucose, 0.15% (w/v) bicarbonate, 0.1 mg/ml streptomycin, 100 U/ml penicillin, 2 mM glutamax and 0.1 mM trolox.

The cultures were kept at 35°C during the imaging procedure. Laser intensity and detector gain were initially set to keep the fluorescent signal in a dynamic range throughout the experiment and were kept constant in all experiments. Of each culture a z-stack was recorded before treatment (day 0) with Δz = 6.3 μm at ideal Nyquist rate and an optical zoom of 0.5. After imaging, cultures were returned to the incubator and treatment was started. The imaging procedure was repeated after 3 days of treatment (day 3) following the same experimental protocol with the exact same imaging parameters. Confocal image stacks were stored as .czi files.

### Immunostaining and imaging

Tissue cultures were fixed in a solution of 4% (w/v) paraformaldehyde (PFA) in PBS (0.1 M, pH 7.4) and 4% (w/v) sucrose for 1 h, followed by 2% PFA and 30% sucrose in PBS (0.1 M, pH 7.4) overnight. Cryostat sections (30 μm; 3050CM S, Leica) of fixed slice cultures were prepared and stained with antibodies against Iba1 (rabbit, 1:1000; Fujifilm Wako #019-19741) and GFAP (mouse, 1:1000; Sigma-Aldrich #G3893). Slices were incubated in appropriate Alexa Fluor^®^ dye-conjugated secondary antibodies (donkey, 1:1000; invitrogen #A-21206). For post-hoc staining, slices were incubated with Alexa Fluor^®^ 488-conjugated streptavidin (1:1000; invitrogen #S32354). TO-PRO^®^-3 (1:5000 in PBS for 10 min; invitrogen #T3605) nuclear stain or DAPI nuclear stain (1:5000 in PBS for 20 minutes; Thermo Scientific #62248) was used to visualize cytoarchitecture. Sections were washed, transferred on glass slides and mounted for visualization with anti-fading mounting medium (Agilent #S302380-2). Confocal images were acquired using a Nikon Eclipse C1si laser-scanning microscope equipped with a 20x (NA 0.75; Nikon) and a 40x oil-immersion objective (NA 1.3; Nikon).

### Propidium iodide staining

Tissue cultures were incubated with propidium iodide (PI, 5 μg/ml; invitrogen #P3566) for 2 h, washed in PBS (0.1 M, pH 7.4) and fixed as described above. Cultures treated for 4 h with NMDA (50 μg/ml; Tocris #0114) and cultures incubated for 2 h with PI after fixation (referred to as ‘post-fixation’) served as positive controls in these experiments. Cell nuclei were stained with TO-PRO^®^-3 (1:5000 in PBS for 10 min; invitrogen #T3605). Cultures were mounted on microscope slides and confocal images were acquired as described above.

### Quantitative reverse transcription PCR (RT-qPCR)

Tissue cultures were transferred into RLT-buffer (QIAGEN) and RNA from individual cultures was isolated according to the manufacturer’s instructions (RNeasy Plus Micro Kit, QIAGEN #74034). Purified RNA was consecutively reverse transcribed (RevertAid RT Kit, Thermo Scientific #K1691). cDNA was diluted in water to a final concentration of 4 ng/μl.

For mRNA correlations with the cytokine detection assay, cDNA preamplification was performed according to the manufacturer’s instruction (TaqMan™ PreAmp Master Mix, Applied Biosystems #4391128) with 25 ng cDNA input and 14 preamplification cycles.

RT-qPCR was performed using a C1000 Touch Thermal Cycler (BIO-RAD) and the CFX 384 Real-Time PCR system (BIO-RAD). 18 ng target cDNA diluted in TaqMan Gene Expression Master Mix (Applied Biosystems #4369016) was amplified using standard TaqMan gene expression assays (Applied Biosystems; Assay-IDs: *Gapdh*: Mm99999915_g1; *Tnf*: Mm00443258_m1; *Il1b*: Mm00434228_m1; *Il6*: Mm00446190_m1; *Il10*: Mm01288386_m1; *Ifng*: Mm01168134_m1). A standard RT-qPCR protocol was used: 1 cycle of 50°C for 2 min, 1 cycle of 95 °C for 10 min, 40 cycles of 95°C for 15 s and 60°C for 1 min. Data were exported and stored on a computer as .pcrd-files.

### Cytokine detection assay

For cytokine detection assay, entorhino-hippocampal tissue cultures were prepared as described above. After preparation, slices were placed on polycarbonate membranes for 24-well plates (two slices per membrane; Thermo Scientific #140620). Cultures were allowed to mature for at least 18 days before experimental assessment. Incubation medium from vehicle-only and LPS-treated tissue cultures was collected, frozen immediately in liquid nitrogen, and stored at −80°C. For detection of TNFα, IFNγ, IL1β, IL6, and IL10 a V-Plex Proinflammatory Panel 1 (mouse) Kit Plus (Mesoscale Discovery #K15048G) was used. Cultures were incubated for three days with vehicle-only, LPS or LPS + IL10 in a 24-well plate as described above, and the incubation medium was collected consecutively. To match gene expression with protein content in the incubation medium, cultures were harvested and processed for RT-qPCR analysis. The collected incubation medium was diluted 1:1 in diluent provided with the kit. Protein detection was performed according to manufacturer’s instructions. A pre-coated plate with capture antibodies on defined spots was incubated with the diluted samples for 2 h. After washing, samples were incubated for 2 h with a solution containing electrochemiluminescent MSD SULFO-TAG detection antibodies (Mesoscale Discovery; Antibodies: Anti-ms IFNγ Antibody #D22QO, Anti-ms IL1β Antibody #D22QP, Anti-ms IL6 Antibody #D22QX, Anti-ms IL10 Antibody, #D22QU, Anti-ms TNFα Antibody #D22QW, Anti-ms IL12p70 Antibody #D22QV). After washing, samples were measured with a MESO QuickPlex SQ 120 instrument (Mesoscale Discovery). The respective protein concentrations were determined using the MSD DISCOVERY WORKBENCH software (Mesoscale Discovery).

### Repetitive magnetic stimulation

Filter membranes containing tissue cultures were transferred to a 35 mm Petri Dish filled with preheated (35°C) standard extracellular solution containing (in mM) 129 NaCl, 4 KCl, 1 MgCl_2_, 2 CaCl_2_, 4.2 glucose, 10 HEPES, 0.1 mg/ml streptomycin, 100 U/ml penicillin, pH 7.4 with KOH, 380-390 mOsm with sucrose. Cultures were stimulated using a Magstim Rapid^2^ stimulator (Magstim Company, UK) that is connected to a Double AirFilm^®^ Coil (coil parameters according to manufacturer’s description: average inductance = 12 μH; pulse rise time approximately 80 μs; pulse duration = 0.5 ms, biphasic; Magstim Company, UK). Cultures were oriented (~ 1 cm beneath the center of the coil) in a way that the induced electric field was parallel to the dendritic tree of CA1 pyramidal neurons. The stimulation protocol consists of 900 pulses at 10 Hz (at 50% of maximum stimulator output). After stimulation, cultures were immediately transferred into culturing medium and kept in the incubator for at least 2 h before experimental assessment. Age- and time-matched control cultures were not stimulated but otherwise treated in the same way.

### Whole-cell patch-clamp recordings

Whole-cell patch-clamp recordings from CA1 pyramidal cells of tissue cultures were carried out at 35°C as described previously (3-7 cells per culture). The bath solution contained (in mM) 126 NaCl, 2.5 KCl, 26 NaHCO_3_, 1.25 NaH_2_PO_4_, 2 CaCl_2_, 2 MgCl_2_, 10 glucose. Patch pipettes contained (in mM) 126 K-gluconate, 10 HEPES, 4 KCl, 4 ATP-Mg, 0.3 GTP-Na_2_, 10 PO-Creatine, 0.3% (w/v) biocytin (pH 7.25 with KOH, 290 mOsm with sucrose), having a tip resistance of 4-6 MΩ. In some experiments Alexa 488 (10 μM; invitrogen #A10436) was added to the internal solution to visualize cellular morphology during electrophysiological assessment. Spontaneous excitatory postsynaptic currents (sEPSCs) were recorded at a holding potential of −60 mV. Series resistance was monitored before and after each recording and recordings were discarded if the series resistance reached ≥ 30 MΩ. For mEPSC recordings, D-APV (10 μM; Abcam #ab120003) and TTX (0.5 μM; Biotrend #18660-81-6) were added to the external solution. For recording of intrinsic cellular properties in current-clamp mode, pipette capacitance of 2.0 pF was corrected and series resistance was compensated using the automated bridge balance tool of the multiclamp commander. IV-curves were generated by injecting 1s square pulse currents starting at −100 pA and increasing in 10 pA steps until +500 pA current injection was reached (sweep duration: 2 seconds).

### Quantification and statistics

RT-qPCR data were analyzed with the Bio-Rad CFX Maestro 1.0 software package using the ΔΔCt method with *Gapdh* as reference gene. Values were normalized to the mean value of the respective vehicle-treated control group.

Colocalization analysis of GFP and Iba1/GFAP was assessed using the ‘coloc2’ plugin of Fiji image processing package (available at https://fiji.sc/; [34]) and Pearson’s correlation coefficients (also referred to as Pearson’s r) were obtained. As a control, Pearson’s r values were calculated from the same set of images in which the Iba1 channel was rotated by 180°.

Mesoscale cytokine detection assay was analyzed using the MSD DISCOVERY WORKBENCH 4.0 platform.

For PI stainings, fluorescence intensity was analyzed in CA1, *stratum pyramidale*. Values were normalized to the mean fluorescence intensity in the vehicle-only treated group.

Confocal image stacks of heterozygous *C57BL/6-Tg(TNFa-eGFP)* cultures were processed and analyzed using the Fiji image processing package. Of each stack a maximum intensity projection followed by background subtraction (rolling ball radius: 50 px) was performed. Culture area was manually defined as a Region of Interest (ROI) and the mean fluorescence intensity of the ROI was calculated. Mean fluorescence intensity of the culture area was normalized to the respective values at day 0 in the same culture.

Single cell recordings were analyzed off-line using Clampfit 11 of the pClamp11 software package (Molecular Devices). sEPSC properties were analyzed using the automated template search tool for event detection. Intrinsic cellular properties were analyzed using Clampfit 11. Input resistance was calculated for the injection of −100 pA current at a time frame of 200 ms with maximum distance to the Sag-current.

Statistical comparisons were performed using non-parametric tests (Mann-Whitney test, Kruskal-Wallis test followed by Dunn’s multiple comparisons). In culture comparison in time-lapse imaging experiments was performed using the Wilcoxon matched-pairs signed rank test. Statistical analysis of cytokine detection assay depicted in figure 5 was performed using a one-way ANOVA test followed by Holm Sidak’s multiple comparison test. p values smaller 0.05 were considered a significant difference. In the text and figures, values represent mean ± standard error of the mean (s.e.m.). * p < 0.05, ** p < 0.01, *** p < 0.001 and not significant differences are indicated by ‘ns’. U values are reported in the figure legends for significant results only.

### Digital illustrations

Figures were prepared using the Photoshop graphics software (Adobe, San Jose, CA, USA). Image brightness and contrast were adjusted.

## RESULTS

### rMS-induced synaptic plasticity is impaired in LPS-treated tissue cultures

The effect of 10 Hz rMS on excitatory synaptic strength was tested in entorhino-hippocampal tissue cultures (≥ 18 days in vitro; Figure 1A, B) prepared from *C57BL/6* mice of both sexes. Tissue cultures were treated with 1 μg/ml of LPS or vehicle-only for three days before the experimental assessment, and AMPA receptor-mediated miniature excitatory postsynaptic currents (mEPSCs) were recorded from individual CA1 pyramidal neurons in whole-cell configuration after the treatment period (Figure 1C-D). Postsynaptic strength was reflected in the amplitude of inward current responses, which are evoked by the stochastic release of glutamate from presynaptic terminals (Figure 1C). Consistent with our previous work [19; 21], increased mEPSC amplitudes were observed 2-4 h following 10 Hz rMS in vehicle-only tissue cultures. Conversely, CA1 neurons of tissue cultures that had been treated with LPS (1 μg/ml, 3 days) prior to magnetic stimulation failed to potentiate their excitatory synapses following rMS (Figure 1D). Importantly, the baseline mEPSC amplitudes of vehicle-only and LPS-treated cultures were not significantly different (Figure 1D). These results demonstrate that LPS administered directly to brain tissue impairs the ability of neurons to express rMS-induced synaptic plasticity without affecting excitatory synaptic strength.

**Figure 1:**
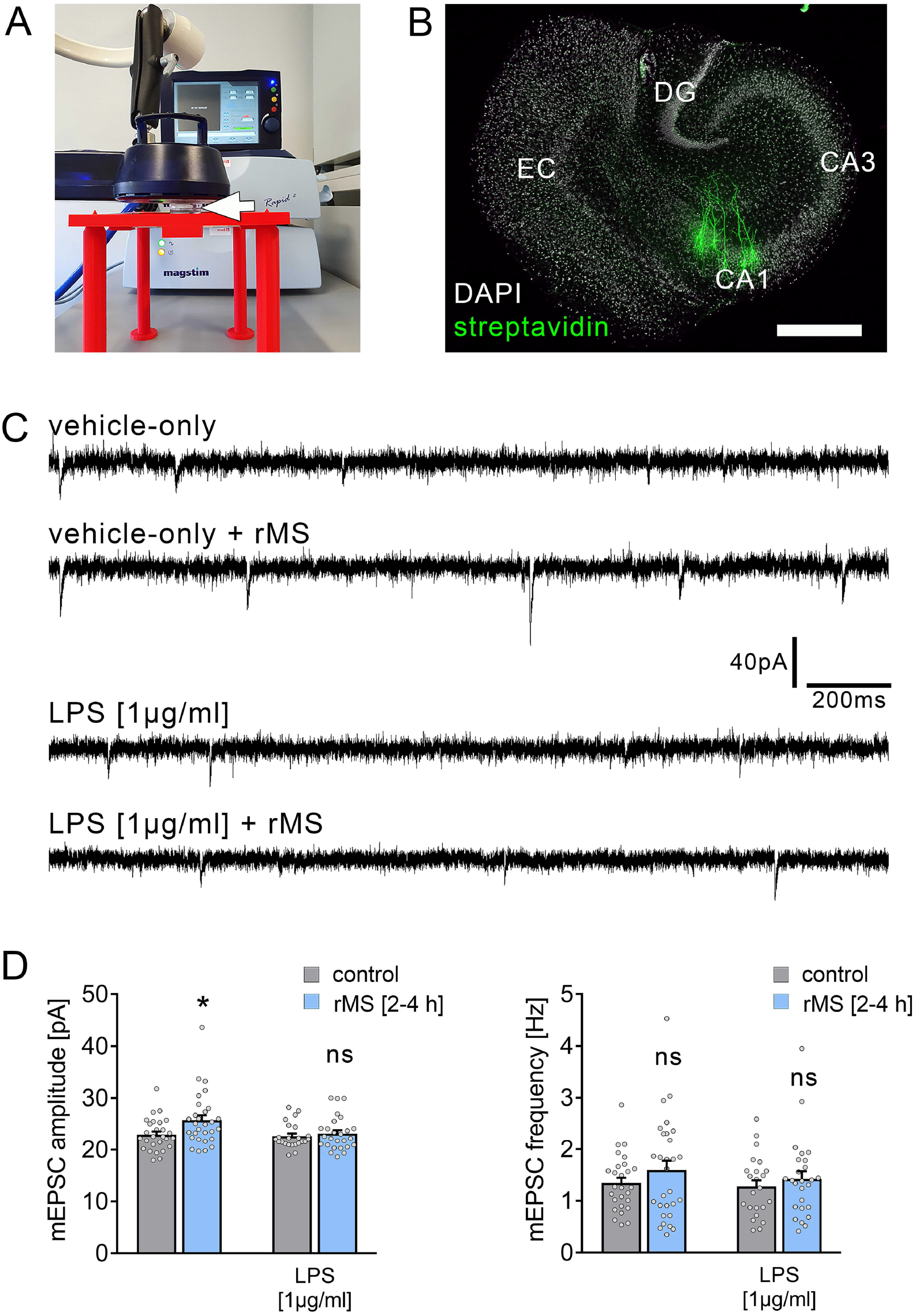
Synaptic plasticity induced by repetitive magnetic stimulation is impaired in entorhino-hippocampal tissue cultures exposed to Lipopolysaccharide. **(A)** Repetitive magnetic stimulation of organotypic entorhino-hippocampal tissue cultures was performed using a Magstim Rapid^2^ stimulator equipped with an AirFilm^®^ stimulation coil. Cultures were placed in a Petri dish below the center of the coil (white arrow) on a custom-designed 3D-printed stage. Distance to coil and orientation of tissue cultures within the electromagnetic field was kept constant in all experiments. A 10 Hz rMS protocol consisting 900 pulses was used to probe excitatory synaptic plasticity. **(B)** Visualization of CA1 pyramidal neurons using Alexa Fluor 488-labeled streptavidin. Scale bar: 500 μm. EC, entorhinal cortex; DG, dentate gyrus. **(C, D)** Representative traces and group data of miniature excitatory postsynaptic currents (mEPSC) recorded from individual CA1 pyramidal neurons 2-4 h after stimulation in the respective groups. Tissue cultures that had been treated with LPS (1 μg/ml, 3 days) prior to magnetic stimulation fail to potentiate their excitatory synapses (vehicle-only: n_control_ = 27 cells, n_rMS [2-4h]_ = 29 cells; Mann-Whitney test, U = 251; LPS: n_control_ = 23 cells, n_rMS [2-4h]_ = 26 cells; Mann-Whitney test; 4 individual rounds, one value excluded in the rMS/vehicle-only group with 65.25 pA/ 3.17 Hz). Individual data points are indicated by gray dots. Values represent mean ± s.e.m. (* p < 0.05; ns, not significant difference). U values are provided for significant results only.

To corroborate these findings, LTP experiments utilizing local electric stimulation of Schaffer collateral-CA1 synapses were carried out in acute slice preparations that had been treated with LPS (1 μg/ml) prior to LTP induction (1 s, 100 Hz; Figure S1). The ability of CA1 pyramidal neurons to express LTP was altered in these experiments, while no changes in baseline synaptic transmission were observed following the application of LPS (at 15 min for ~ 6 h, i.e., prior to LTP induction in the LPS-treated group; Figure S1). These findings agree with our previous *in vivo* work [23], thus validating the use of *in vitro* LPS-exposure as a method to study neuroinflammation-associated alterations in synaptic plasticity.

Together, these experiments demonstrate that both rMS-induced synaptic plasticity and classic electric tetanus-induced LTP are altered by direct exposure to LPS.

### LPS-induced neural toxicity does not account for alterations in rMS-induced synaptic plasticity

Propidium iodide (PI) stainings were used to assess cell viability in our experimental setting. PI is a cell membrane impermeant fluorescent molecule that binds to DNA. Hence, PI can be used as a marker for membrane integrity when applied to living tissue [35].

To test for LPS-induced effects on cell viability in organotypic tissue cultures, another set of cultures was treated with LPS (1 μg/ml, 3 days) and vehicle-only. Formaldehyde fixation, which compromises membrane integrity, and short-term NMDA treatment (50 μM, 4 h) served as positive controls in this series of experiments. Figure 2 shows that the PI-signal was comparably low in the control and LPS-treated groups, while a significant increase in PI-signal was evident in the NMDA-treated group, and almost all nuclei were PI-positive following fixation (Figure 2A-C).

**Figure 2:**
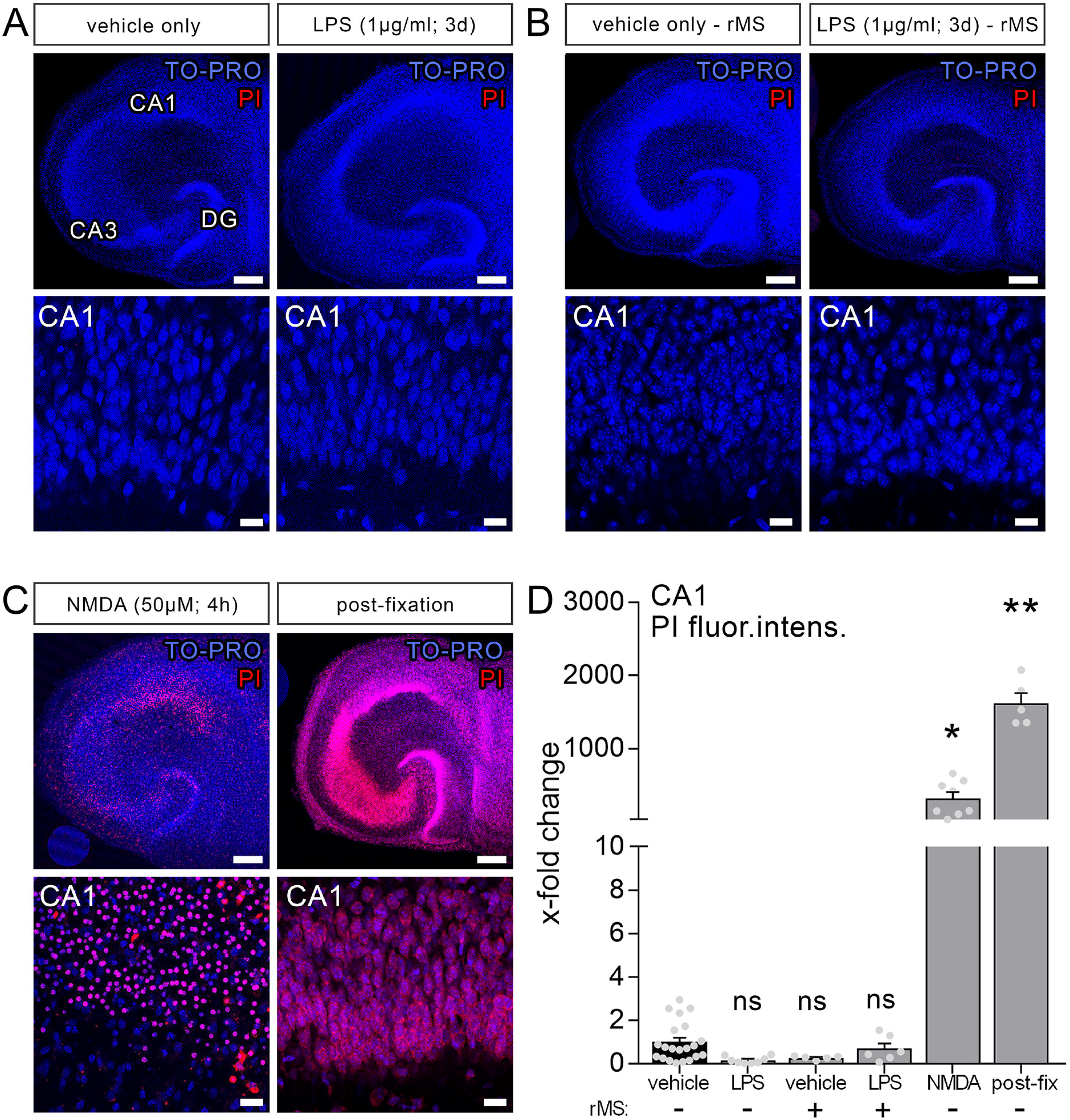
Lipopolysaccharide and/or repetitive magnetic stimulation do not affect cell viability. **(A-C)** Representative examples of propidium iodide staining (PI, red) in the various experimental conditions (TO-PRO nuclear stain, blue). (A) No effect on membrane integrity is observed following *in vitro* LPS exposure (1 μg/ml, 3 days) as compared to age- and time-matched vehicle-only treated controls. (B) No signs of cell death are detected 2-4 h after rMS, both in vehicle-only and LPS-treated tissue cultures. (C) NMDA treatment (50 μM, 4 h) and formalin fixation served as positive controls in these experiments. Scale bars: 200 μm and 20 μm. **(D)** Summary graph and combined analysis of changes in propidium iodide fluorescence intensity in area CA1 under the indicated experimental conditions, respectively (n_vehicle-only_ = 21 cultures, n_LPS_ = 8 cultures, n_rMS_ = 5 cultures, n_LPS+rMS_ = 6 cultures, n_NMDA_ = 8 cultures, n_post-fix_ = 5 cultures; Kruskal-Wallis test followed by Dunn’s multiple comparisons). Individual data points are indicated by gray dots. Values represent mean ± s.e.m. (* p < 0.05, ** p < 0.01; ns, not significant difference).

Considering the clinical relevance of non-invasive magnetic stimulation, i.e., TMS, we also tested for the effects of 10 Hz rMS. No apparent negative effects on cell viability (2-4 h after stimulation) were detected in the vehicle-only controls, which exhibited rMS-induced synaptic plasticity, or in the LPS-treated cultures, in which no increase in excitatory synaptic strength following 10 Hz rMS was evident (Figure 2B). According to the above results (Figure 2D), we conclude that neither LPS nor 10 Hz rMS significantly affects cell viability. This finding corroborates previous work in which no negative or even neuroprotective effects of rTMS have been reported [36; 37; 38].

### LPS triggers microglial TNFα expression in organotypic tissue cultures

To further characterize the effects of LPS in our experimental setting, we used entorhino-hippocampal tissue cultures prepared from a novel transgenic reporter mouse line [*C57BL/6-Tg(TNFa-eGFP)*], which expresses eGFP under the control of the *Tnfa* promoter (Figure 3A, B). Cultures prepared from these mice were treated with LPS (1 μg/ml, 3 days); baseline fluorescence intensity was measured prior to treatment (day 0) and at the end of the treatment period (day 3) using live-cell microscopy. A significant increase in the eGFP-fluorescence was detected in the LPS-treated group as compared to the vehicle-only controls, which did not show significant changes in eGFP expression (Figure 3C, D). The validity of this experimental approach was further supported by correlated changes between eGFP-fluorescence and RT-qPCR analysis for *Tnfa*-mRNA in the same set of tissue cultures (Figure 3D).

**Figure 3:**
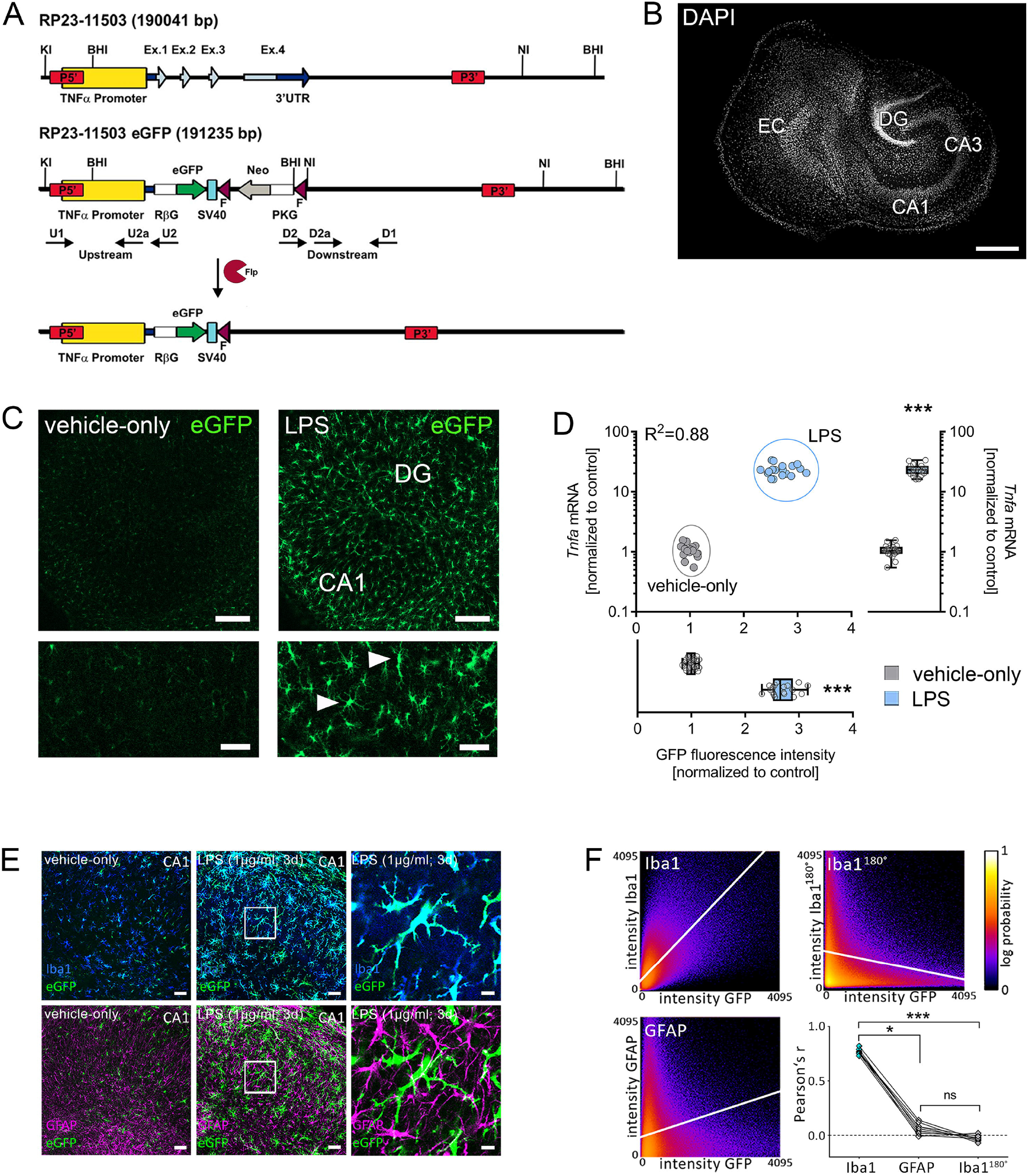
Microglia are a major source of tumor necrosis factor α (TNFα) in tissue cultures exposed to Lipopolysaccharide. **(A, B)** Schematic representation of the *Tg(TNFa-eGFP)* reporter transgenic construct. The *Tnfa* locus of BAC RP23-115O3 containing the *Tnfa* promoter as well as the four exons (Ex. 1 to 4) are shown in the upper part. In the recombinant eGFP BAC (middle part), the *Tnfa* coding sequence and the 3’ untranslated region (3’UTR) were replaced by eGFP and neomycin (Neo) coding cassettes. After flippase (Flp) mediated intra-chromosomal recombination, the neo cassette is excised, leaving the Rabbit β-globin intron (RβG)-eGFP fusion gene under the control of the *Tnfa* promoter, as illustrated in the lower part. SV40, SV40 early mRNA polyadenylation; F, flippase recognition target; PKG, phosphoglycerate kinase. A tissue culture stained with DAPI nuclear staining is shown to visualize cytoarchitecture. Scale bar: 400 μm. **(C)** Examples of tissue cultures prepared from *C57BL/6-Tg(TNFa-eGFP)* mice after 3 days of treatment with 1μg/ml LPS (right side) and vehicle-only (left side). A considerable increase in the eGFP signal is observed in the LPS group (smaller sections with higher magnification at the bottom). Scale bars: 200 μm and 75 μm. **(D)** Quantification of the eGFP signal and the corresponding *Tnfa*-mRNA content of individual cultures in vehicle-only (grey dots) and LPS-treated (blue dots) *C57BL/6-Tg(TNFa-eGFP*) organotypic tissue cultures (n = 18 cultures per group and experimental condition; Mann-Whitney test, UGFP-intensity = 0, U*Tnf*-mRNA = 0; Boxplot diagrams indicate the level of significance and distribution of data points within the groups). **(E)** Examples of *C57BL/6-Tg(TNFa-eGFP)* tissue cultures treated with vehicle-only or LPS stained for the astrocytic marker GFAP and the microglial marker Iba1. Scale bars: 50 μm and 10 μm. **(F)** Quantitative assessment of colocalization shows colocalization between the GFP- and the Iba1-signal, but not the GFP- and the GFAP-signal. The same set of images in which the Iba1-signal is rotated 180° served as a control for this analysis (n = 9 cultures per group; Kruskal-Wallis test followed by Dunn’s multiple comparisons). Individual data points are indicated by single dots. Values represent mean ± s.e.m. (* p < 0.05, *** p < 0.001; ns, not significant difference).

The majority of eGFP-expressing cells were Iba1 and not GFAP positive as shown by immunostaining of tissue cultures (Figure 3E, F). Careful examination of Iba1 and GFAP negative areas in CA1 *stratum pyramidale*, i.e., the layer in which the cell bodies of the recorded CA1 neurons are found, revealed no evidence of neuronal TNFα expression. Although these experiments did not exclude astrocytic and/or neuronal sources of TNFα, our data demonstrate that microglia are a major source of TNFα during LPS-induced neuroinflammation. This observation is in line with previous work that has demonstrated the presence of toll-like receptor 4 (TLR4) on microglia, i.e., the main receptor targeted by LPS [39; 40].

### IL10 attenuates LPS-induced TNFα expression in tissue cultures prepared from TNFα-reporter mice

Next, we tested for the effects of the anti-inflammatory cytokine IL10 in LPS-treated tissue cultures (Figure 4) [41; 42]. Tissue cultures prepared from TNFα-reporter mice were again exposed to LPS (1 μg/ml, 3 days), and changes in eGFP-fluorescence were monitored using live-cell microscopy (Figure 4A). While a robust increase in eGFP-fluorescence was detected in the LPS-treated cultures, IL10 (10 ng/ml) attenuated the increase in eGFP-fluorescence in LPS-treated tissue cultures (Figure 4A-C). Similar results were obtained, when RT-qPCR analysis for *Tnfa*-mRNA was carried out at the end of each experiment in the same set of tissue cultures (Figure 4D). The results demonstrate the anti-inflammatory effects of IL10 in LPS-treated tissue cultures, which is in line with recent reports [28]. Thus, we conclude that tissue cultures prepared from our new TNFα-reporter mouse line are suitable tools for studying neural inflammation and anti-inflammatory intervention.

**Figure 4:**
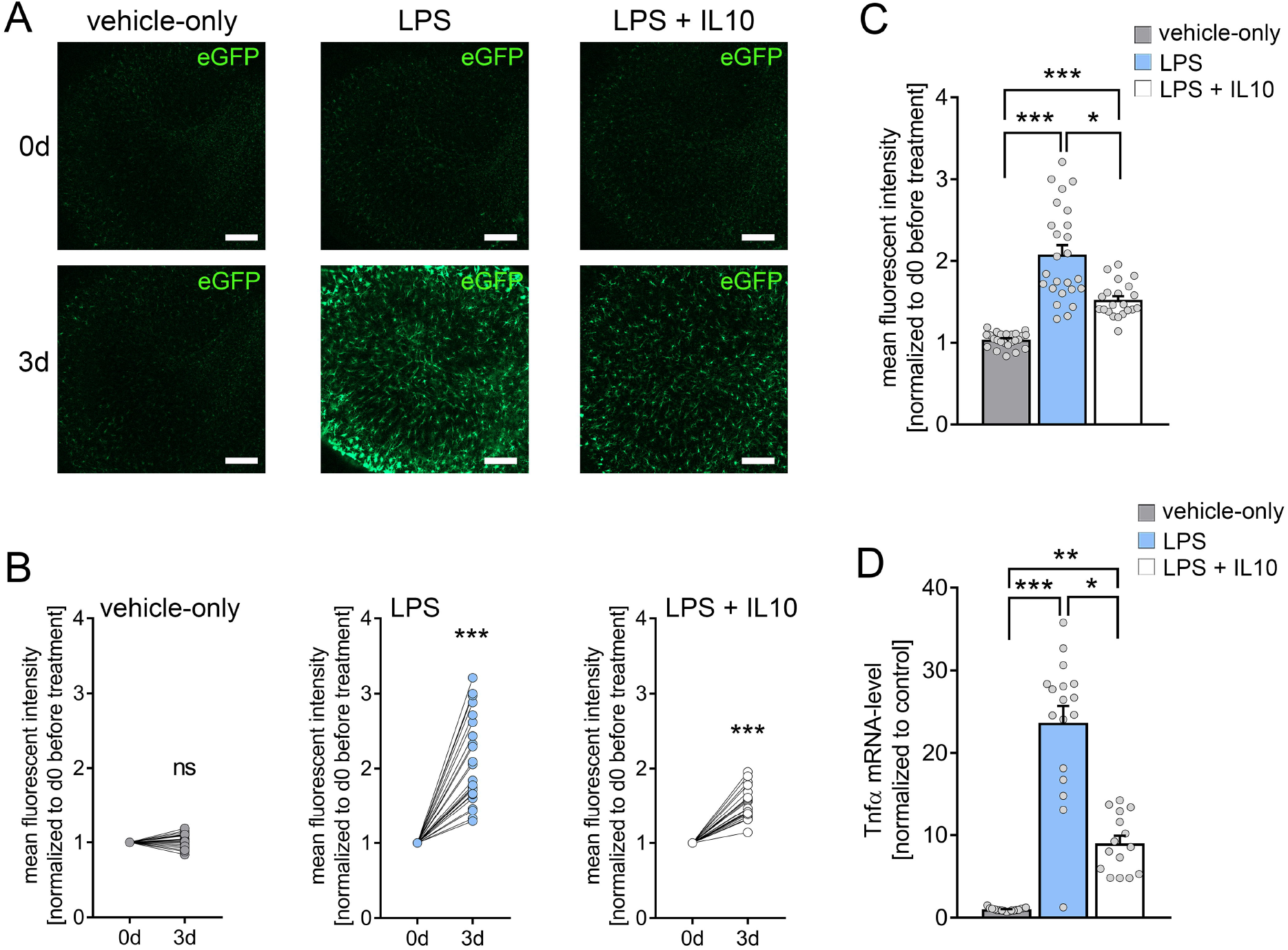
Interleukin 10 (IL10) attenuates *in vitro* LPS-induced TNFα expression. **(A)** Examples of time-lapse imaging of *C57BL/6-Tg(TNFa-eGFP)* tissue cultures before treatment (0 d, row at the top) and 3 days after treatment with vehicle-only, LPS (1 μg/ml) or LPS (1μg/ml) + IL10 (10 ng/ml), respectively (3 d, row at the bottom). Scale bars: 200 μm. **(B, C)** Group data of eGFP fluorescence intensities before the treatment (0 d) and after 3 days in the respective groups (n_vehicle-only_ = 24, n_LPS_ = 25, n_LPS+IL10_ = 21 cultures per group; Wilcoxon matched-pairs signed rank test). IL10 restrains LPS-induced microglia TNFa expression (Kruskal-Wallis test followed by Dunn’s multiple comparisons). **(D)** Changes in eGFP fluorescence intensity correspond to *Tnfa*-mRNA levels in the respective groups (n_vehicle-only_ = 17, n_LPS_ = 17, n_LPS+IL10_ = 15 cultures per group; Kruskal-Wallis test followed by Dunn’s multiple comparisons test). Individual data points are indicated by colored dots. Values represent mean ± s.e.m. (* p < 0.05, ** p < 0.01, *** p < 0.001; ns, not significant difference).

### The effects of LPS and IL10 on TNFα, IL6, IL1β, IL10 and IFNγ expression in organotypic tissue cultures

A sandwich-ELISA approach was employed to confirm the above results at the protein level. The culturing medium of the vehicle-only controls, LPS, and LPS + IL10 treated tissue cultures prepared from C57BL/6 mice was analyzed for TNFα. Tissue cultures were subjected to further qPCR analysis. Figure 5A shows that while TNFα was barely evident in the medium of vehicle-only treated controls, the LPS-treated group demonstrated a significant increase in TNFα. Consistent with our mRNA results (all Ct-values are provided in supplemental Table S2), IL10 restrained a significant increase in TNFα protein levels upon exposure to LPS (Figure 5A, c.f. Figure 4D; for values see Table S3). We also tested for the effects of LPS and LPS + IL10 on the pro-inflammatory cytokines IL6, IL1β, and IFNγ and found similar results (Figure 5B-D). Specifically, IL6 protein levels increased significantly in the LPS-treated group and this increase was attenuated in the presence of IL10. The anti-inflammatory effect of IL10 was evident in all pro-inflammatory cytokines tested, i.e., IL6, IL1β, and IFNγ. Interestingly, the change in *Ifng-*mRNA levels did not reflect altered IFNγ protein concentrations in the medium. While LPS triggered a reduction in *Ifng*-mRNA, a slight but highly significant increase in IFNγ was detected (Figure 5D). From these results, we conclude that organotypic tissue cultures are suitable tools to study LPS-induced neural inflammation and anti-inflammatory treatment strategies, e.g., IL10 administration.

**Figure 5:**
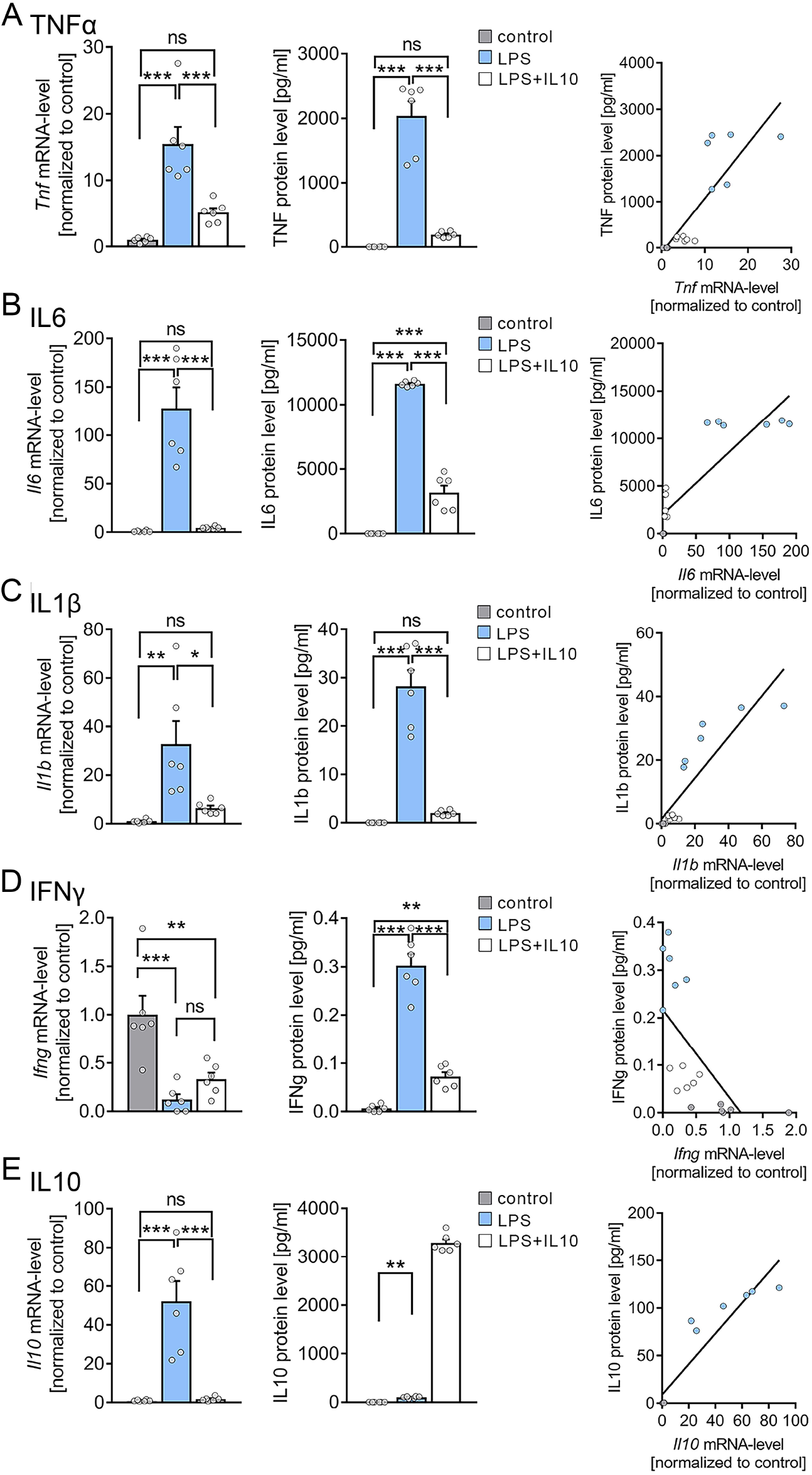
Expression profiles of cytokines in entorhino-hippocampal tissue cultures exposed to Lipopolysaccharide and interleukin 10 (IL10). **(A-D)** Group data and correlations of mRNA and protein levels in the incubation medium of (A) tumor necrosis factor α (TNFα), (B) interleukin 6 (IL6), (C) interleukin 1β (IL1β), and interferon γ (IFNγ) in vehicle-only, LPS (1 μg/ml, 3 d) or LPS (1 μg/ml, 3 d) + IL10 (10 ng/ml, 3 d) treated cultures, respectively. **(E)** Group data of mRNA and protein levels of IL10 in the respective groups (one-way ANOVA followed by Holm-Sidak’s multiple comparison test, Mann-Whitney test for statistical comparison of IL10 levels (U = 0); linear regression fit indicated by black line; values for IL10 protein level in the LPS + IL10 treated group were excluded from the fitting procedure; n = 6 cultures or culturing medium respectively for each experimental condition). Individual data points are indicated by colored dots. Values represent mean ± s.e.m. (* p < 0.05, ** p < 0.01, *** p < 0.001; ns, not significant difference). U values are provided for significant results only.

### IL10 levels in organotypic tissue cultures

Next, we examined changes in endogenous *Il10*-mRNA and IL10 protein levels (Figure 5E). A significant increase in *Il10*-mRNA levels was detected in the LPS group, which was prevented by the application of recombinant IL10. Consistent with these findings, IL10 protein levels significantly increased in the medium of LPS-treated tissue cultures, and the exogenously applied recombinant IL10 was readily detected in the LPS + IL10 group (Figure 5E). We conclude, therefore, that an endogenous source of IL10 exists in our tissue cultures. Consistent with a negative-feedback mechanism, this endogenous IL10 source is obstructed by the administration of recombinant IL10. It appears that the endogenous source of IL10 is insufficient to prevent LPS-induced alterations in synaptic plasticity.

### IL10 restores 10 Hz rMS-induced synaptic plasticity in LPS-treated tissue cultures

To address the question of whether IL10 is able to restore rMS-induced synaptic plasticity in LPS-treated tissue cultures, we repeated the experiments described above and stimulated once more tissue cultures with 10 Hz rMS. Basic intrinsic properties (Figure 6A-C) and spontaneous excitatory postsynaptic currents (sEPSCs, Figure 7) were recorded from CA1 pyramidal neurons.

**Figure 6:**
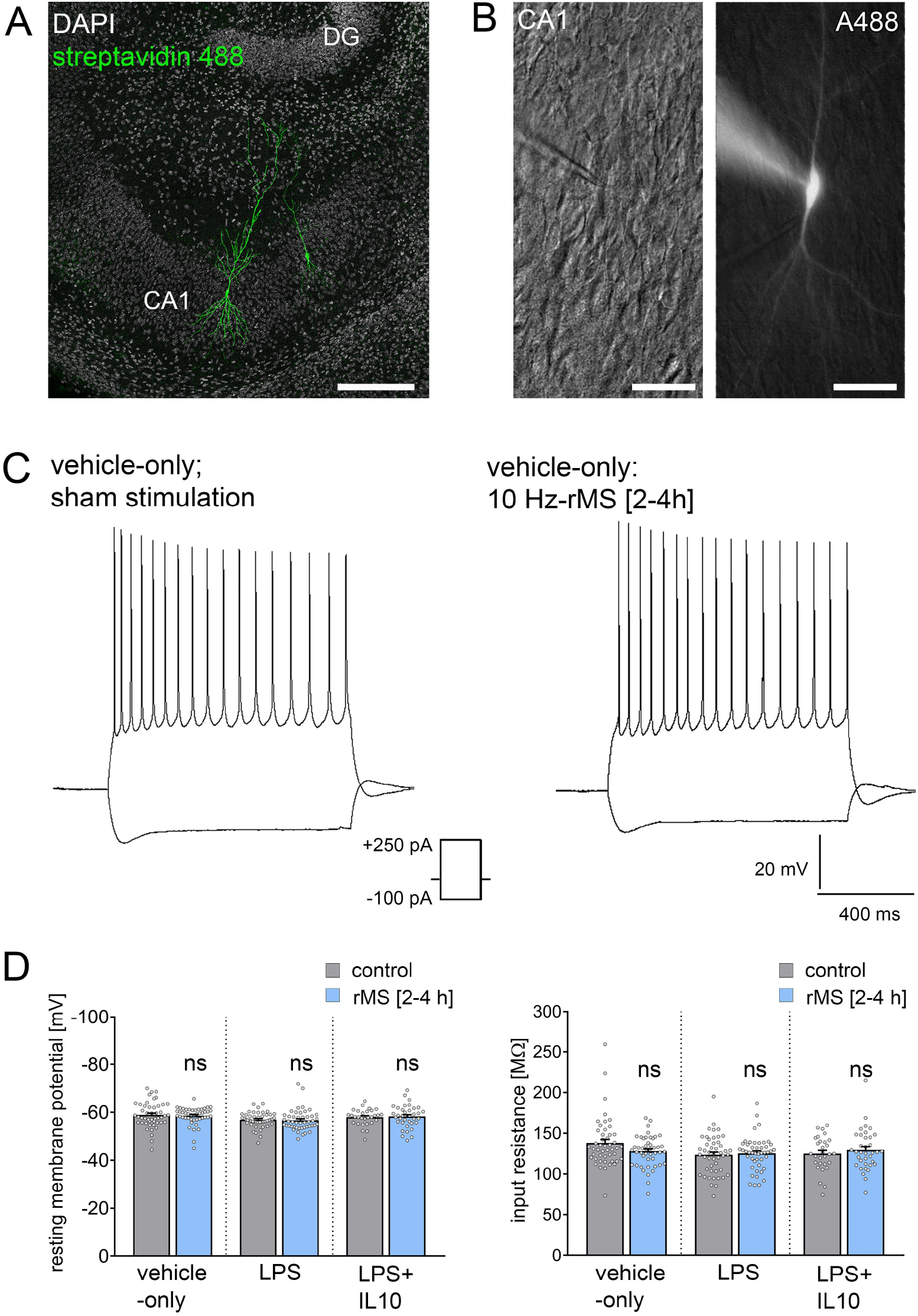
Repetitive magnetic stimulation (rMS) and Lipopolysaccharide have no major impact on intrinsic cellular properties of CA1 pyramidal neurons. **(A, B)** Examples of CA1 pyramidal neurons recorded and filled with Alexa 488 (10 μM). Cytoarchitecture visualized with DAPI in (A), Dot gradient contrast in (B). Scale bar: (A) 200 μm, (B) 50 μm. **(C)** Sample traces and group data of input-output properties in CA1 pyramidal neurons from vehicle-only, LPS (1 μg/ml) or LPS (1μg/ml) + IL10 (10 ng/ml) treated (3 d) tissue cultures, respectively. **(D)** Resting membrane potential and input resistance is not significantly different between the groups (vehicle-only: n_control_ = 47 cells, n_rMS [2-4h]_ = 46 cells; LPS: n_control_ = 47 cells, n_rMS [2-4h]_ = 46 cells; LPS + IL10: n_control_ = 29 cells, n_rMS [2-4h]_ = 36 cells; Mann-Whitney test; one recording excluded in the LPS/rMS-group due to high series resistance during recordings). Individual data points are indicated by colored dots. Values represent mean ± s.e.m. (ns, not significant difference).

**Figure 7:**
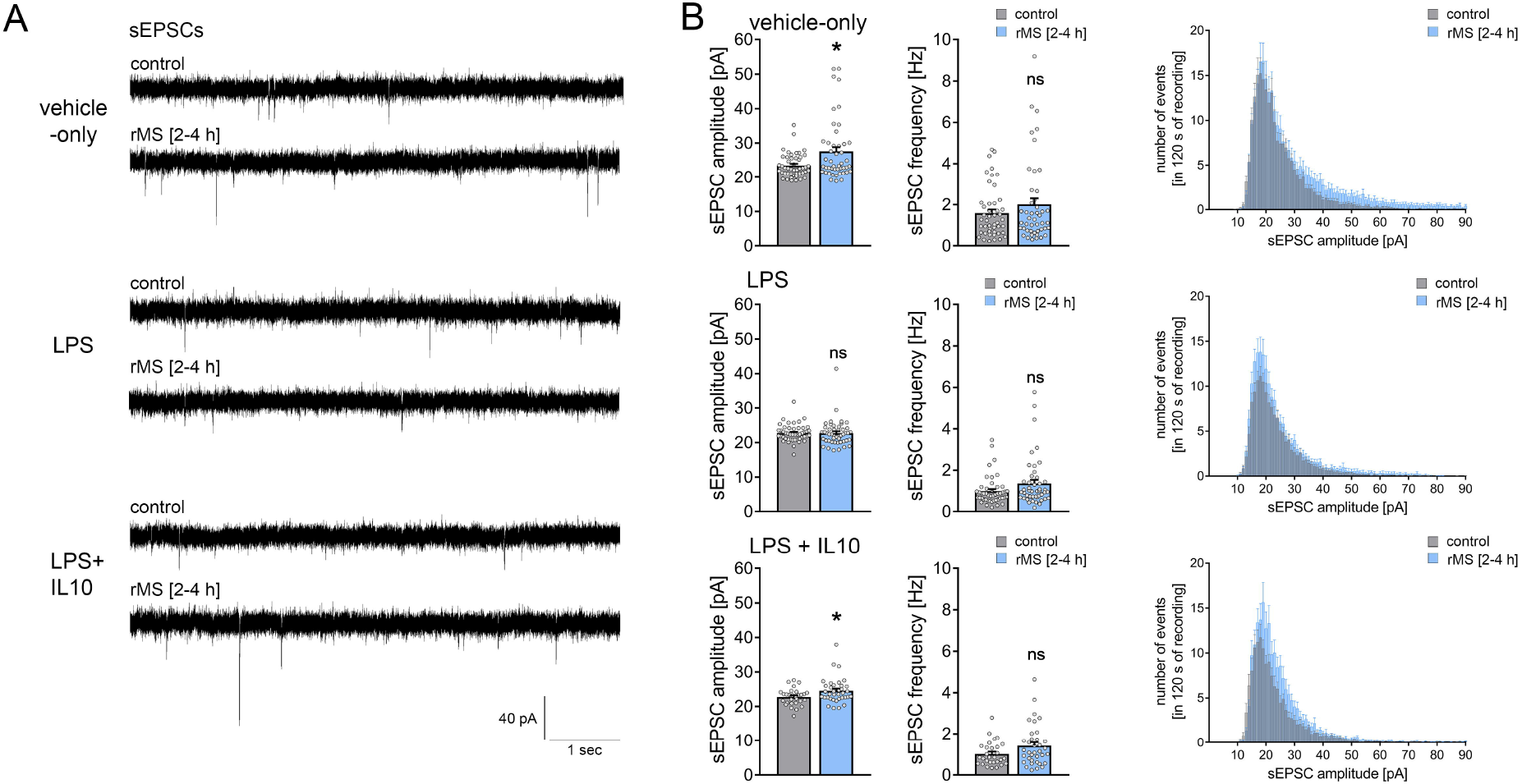
Interleukin 10 (IL10) restores the ability of CA1 pyramidal neurons to express synaptic plasticity in tissue cultures exposed to Lipopolysaccharide.. **(A, B)** Sample traces and group data of spontaneous excitatory postsynaptic currents (sEPSCs) recorded from CA1 pyramidal neurons of vehicle-only, LPS (1 μg/ml) or LPS (1μg/ml) + IL10 (10 ng/ml) treated (3 d) tissue cultures, respectively. 10 Hz repetitive magnetic stimulation (900 pulses) was used to probe excitatory synaptic plasticity (vehicle-only: n_control_ = 47 cells, n_rMS [2-4h]_ = 46 cells; LPS: n_control_ = 47 cells, n_rMS [2-4h]_ = 47 cells; LPS + IL10: n_control_ = 29 cells, n_rMS [2-4h]_ = 36 cells; 4 individual rounds; Mann-Whitney test, vehicle-only: U_sEPSC amplitude_ = 791, LPS + IL10: U_sEPSC amplitude_ = 372). Individual data points are indicated by colored dots. Values represent mean ± s.e.m. (* p < 0.05, ns, not significant difference). U values are provided for significant results only.

Figure 6 shows that there were no major difference in input-output properties and no significant difference in the resting membrane potential or input resistance between stimulated and non-stimulated tissue cultures (Figure 6C, D). Similar results were obtained from tissue cultures treated with LPS (1 μg/ml, 3 days) and LPS (1 μg/ml, 3 days) + IL10 (10 ng/ml, 3 days). We conclude from these results that 10 Hz rMS does not affect the basic functional properties of CA1 pyramidal neurons.

Recordings of AMPA-receptor mediated sEPSCs in the same set of neurons (Figure 7A) confirmed the previously reported rMS-induced potentiation of excitatory neurotransmission (Figure 7B). Likewise, the negative effects of LPS on rMS-induced synaptic plasticity were again observed in these experiments (Figure 7B). Finally, in the LPS + IL10-treated group, a significant increase in sEPSC amplitudes was detected following rMS (Figure 7B). Overall, these results are evidence that the anti-inflammatory cytokine IL10 restores the ability of neurons to express synaptic plasticity probed by 10 Hz rMS in an *in vitro* model of LPS-induced neural inflammation.

## DISCUSSION

A substantial body of research supports the fact that inflammatory cytokines and other immune mediators are capable of affecting synaptic transmission and plasticity [25; 27; 30; 43; 44; 45]. Indeed, bacteremia and sepsis trigger complex responses in the central nervous system including microglia activation and alterations in synaptic plasticity [46; 47]. However, the underlying mechanism between microglia activation and synaptic plasticity under physiological and pathological conditions remains poorly understood (c.f., [48]). The results of the present study support organotypic tissue cultures as tools to study synaptic plasticity under the conditions of neural inflammation and to test for anti-inflammatory and/or plasticity-promoting interventions. Moreover, findings validate the use of tissue cultures prepared from a new transgenic mouse line, i.e., TNFα-reporter mice, and identify microglia as a major source of LPS-induced TNFα and, hence, neural inflammation in our experimental setting. Using this approach, we demonstrate that (i) IL10 restrains LPS-induced neural inflammation *in vitro* and (ii) restores synaptic plasticity probed by rMS in tissue cultures exposed to LPS. These results provide empirical support for the use of rTMS as a diagnostic tool in the context of brain inflammation.

Our previous study demonstrated that the *in vivo* intraperitoneal injection of LPS is accompanied by an increase in brain TNFα levels and alterations in the ability of hippocampal CA1 pyramidal neurons to express excitatory synaptic plasticity [23]. Interestingly, in the absence of an LPS-triggered serum cytokine pulse, i.e., when LPS is administered directly to brain tissue (*in vitro),* similar effects on synaptic plasticity are observed. While baseline synaptic transmission is not affected, the ability to potentiate excitatory neurotransmission is altered in the presence of LPS, both following electric tetanic stimulation and 10 Hz rMS *in vitro*. These findings indicate that LPS can act directly on neural tissue and that it is sufficient to trigger alterations in synaptic plasticity in the CNS. Notably, these effects are not readily explained by LPS-mediated neurotoxicity as shown by the PI stainings in our tissue cultures, and the findings do not support rTMS as affecting cell viability. Yet, the precise role of TLR4-mediated microglia activation [49] warrants further investigation. It remains to be investigated, for example, whether LPS impairs plasticity via microglia activation, which in turn could trigger TNFα expression (and other cytokines), or via additional direct effects on astrocytes and neurons [50; 51].

The effects of LPS on synaptic plasticity concur with the findings of our recent work on the concentration-dependent effects of TNFα in CA1 pyramidal neurons [30]. In this earlier study, we showed that a high concentration of TNFα impairs the expression of excitatory synaptic plasticity, which reflects the LPS effects reported in the present study. It is conceivable, therefore, that TNFα contributed to the detrimental effects of LPS on synaptic plasticity in our experimental setting, which is supported by recent studies on microglial TNFα deficiency [28]. Considering the results of our sandwich-ELISA analysis, future research should extend this exploration to investigate the role of the concentration-dependent effects of IL6, IL1β, and IFNγ on synaptic plasticity modulation.

Regardless of these considerations, we identified microglia as a major source of LPS-induced TNFα in tissue cultures prepared from our new TNFα-reporter mouse line. Changes in eGFP expression were reflected in *Tnfa*-mRNA expression and altered TNFα protein levels in the culturing medium, thus validating the use of TNFα-reporter mice. Furthermore, the anti-inflammatory effects of IL10 were readily detected using live-cell microscopy and quantification of the eGFP signal (for recent work on the effects of IL10 in the central nervous system see [52; 53; 54]). Hence, tissue cultures prepared from these mice are suitable to monitor neural inflammation and anti-inflammatory interventions *in vitro*. Certainly, crossing *C57BL6-Tg(TNFa-eGFP)* mice with other suitable transgenic mouse lines (e.g., HexB-tdTomato; [55]) could be instrumental to better understand the complex role of microglial TNFα in physiological and pathological conditions [56; 57].

In this context, several recent studies have implicated microglia and defective IL10 signaling in various brain disease and lesion models [58; 59; 60; 61]. Further, IL10 expression has been reported for cultured microglia [62]. However, to date, transcriptome analyses of microglia isolated from various *in vivo* disease models have failed to demonstrate IL10 expression in microglia [28; 63; 64; 65]. For example, recovery from spinal cord injury depended on IL10, which was produced by monocytes and not microglia [66]. Considering that cultured microglia lose their characteristic tissue imprints and adopt a gene expression signature that resembles prototype macrophages [67; 68], it could be assumed that microglia are not capable of producing IL10 in a mature (organotypic) environment. Our experiments in mature, i.e., three-week old tissue culture preparations show that LPS triggers endogenous IL10 expression, both at the mRNA and protein level. However, when recombinant IL10 was administered *Il10*-mRNA expression was reduced to control levels. These findings are consistent with a prominent negative feedback mechanism that regulates IL10 expression in organotypic tissue cultures. However, it could be considered that the controversy between studies on microglial IL10 expression may depend, at least in part, on such negative feedback mechanisms and, hence, on the presence of other IL10 sources in the respective experimental settings, e.g., the presence of monocytes, T-cells and other peripheral immune cells, which are absent in three-week old tissue cultures. Future studies should explore the relevance of such negative feedback mechanism and genomic regulatory elements that suppress microglial IL10 production in the presence of IL10 and other restricting factors. We are confident that our organotypic tissue culture approach will be helpful in addressing these and other important questions. Accordingly, testing for the effects of neural activity and brain stimulation in this context will be an interesting avenue for future research. The results of the present study provide robust evidence that non-invasive brain stimulation techniques, i.e., rTMS, are suitable tools for monitoring alterations in synaptic plasticity in brain inflammation and the plasticity restoring effects of IL10.

## DECLARATIONS

### ETHICS STATEMENT

All experimental procedures were performed according to the national animal welfare legislation and approved by the animal welfare committee (V54-19c20/15-F143/37), the local animal welfare officers (Universities of Frankfurt, Mainz and Freiburg) and/or the institutional animal care and use committee (Sheba Medical Center).

### AVAILABILITY OF DATA AND MATERIALS

The datasets generated and/or analyzed during the current study are available from the corresponding author on reasonable request.

### AUTHORS CONTRIBUTION

Roles of authors and contributors have been defined according to ICMJE guidelines and author contributions have been reported according to CRediT taxonomy. Conceptualization, M.L., S.J., N.M. and A.V.; Methodology, M.L., N.M. and A.V.; Formal Analysis, M.L., A.E., P.K. A.S., S.R. and N.M.; Investigation, M.L., A.E., P.K. A.S., S.R., and N.M.; Resources, I.G., N.Y., S.F., A.W. and A.V.; Writing Original Draft, M.L. and A.V.; Visualization, M.L., A.E., and A.V.; Supervision, N.M. and A.V.; Project Administration, M.L., T.D., N.M. and A.V.; Funding Acquisition, M.L., T.D., J.S., N.M. and A.V.

