## Supplementary Material for "Neural inflammation alters synaptic plasticity probed by 10 Hz repetitive magnetic stimulation"

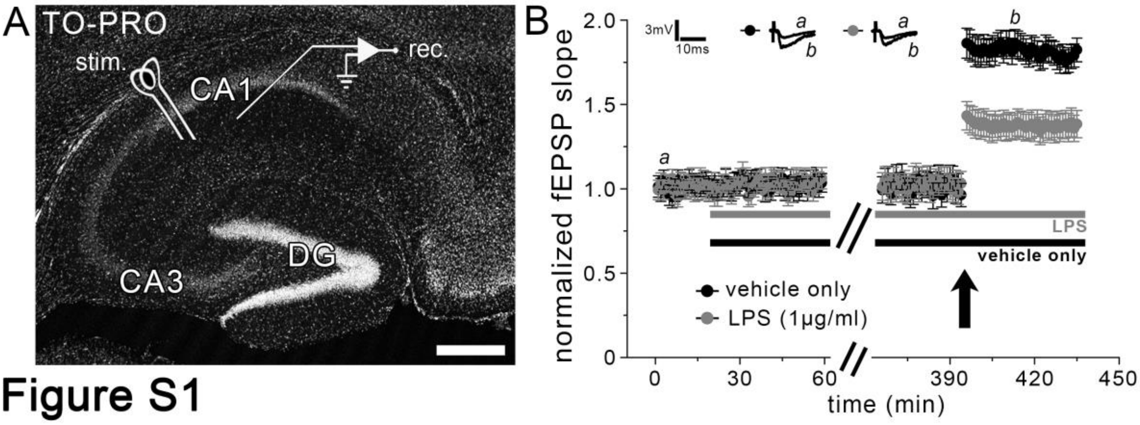


**Figure S1: Long-term potentiation of Schaffer collateral-CA1 synapses is impaired in Lipopolysaccharide-treated acute hippocampal slices.**

**(A, B)** A single 100 Hz electric tetanus (1 s; indicated by arrow in B) is applied to induce LTP at Schaffer collateral-CA1 synapses either in LPS (1 µg/ml) or vehicle-only treated acute hippocampal slices (TO-PRO nuclear stain; Scale bar: 300 µm). While LPS treatment does not affect baseline synaptic transmission, the ability of neurons to express synaptic plasticity is impaired. Representative traces of field excitatory postsynaptic potentials (fEPSP) at indicated times (a, b) are shown on top (n = 12 slices from 4 animals for each group; unpaired two-tailed t-test; $p < 0.001$).

Values represent mean ± s.e.m.

**Table S1: Primers used in the generation of *C57BL/6-Tg(TNFa-eGFP)* mouse line.**

| | Primer | Sequence |
|---|---|---|
| **1a** | RbGLClaIf | gcatatcgatcctgagaacttcagggtga |
| **1b** | RbGL-ERIr | gcatgaattcgccctatagtgagtcgtattaca |
| **2a** | eGFP-ERIf | gcatgaattccaccatggtgagcaagggcga |
| **2b** | eGFP-PAf | ggcatggacgagctgtacaagtaatctagatcataatcagccataccaca |
| **2c** | eGFP-PAr | tgtggtatggctgattatgatctagattacttgtacagctcgtccatgcc |
| **2d** | SV40PA-SalIr | gcatgtcgacttaagatacattgatgagt |
| **3a** | mTNFp-NIBBIf | cagtgcggccgcttcgaagctctaaaagccagccact |
| **3b** | mTNFp-HdIIIr | gcataagcttggtgtcttttctggagggaga |
| **4a** | PGNeoFRT-NheIf | cgatgctagcggggtaaccgaagttcctatactttctag |
| **4b** | PGNeoFRT-SalIr | tggcgtcgactcgcattttgaagttcctattccgaagttcc |

| | | |
|---|---|---|
| **5a** | TNFds-NheIf | gcatgctagcgtgatttctgtcttgggatgaagt |
| **5b** | TNFds-KSr | tgacggtacccggggctcttaagacccacttgct |
| **6a** | U1 | ctaggtcccagacacaaagg |
| **6b** | U2 | gatacaagggacatcttccc |
| **6c** | U2a | aagcttggtgtcttttctggagggag |
| **7a** | D1 | atacctagtcattgccttcc |
| **7b** | D2 | acccggtagaattgacctgc |
| **7c** | D2a | ttcccactctgggaattcc |
| **8a** | RP23Southf | atcttctcaacctggatggg |
| **8b** | RP23Southr | atccagccacccaacccc |
| **9a** | RP23SouthUpf | atccccaccagtggcctc |

| | | |
|---|---|---|
| **9b** | RP23SouthUpr | atctaattctctcgccatctc |
| **10a** | PR23115f (Seq) | ctaggtcccagacacaaagg |
| **10b** | PR23115r (Seq) | atacctagtcattgccttcc |

**Table S2: Ct-values for qPCR analysis in mRNA/protein-correlation analysis.**

| | **vehicle-only** | **LPS** | **LPS + IL10** |
|---|---|---|---|
| ***Tnfa*** | 25.10 ± 0.23 | 20.12 ± 0.15 | 20.77 ± 0.22 |
| ***Il6*** | 31.24 ± 0.32 | 23.19 ± 0.30 | 26.98 ± 0.33 |
| ***Il1b*** | 25.07 ± 0.41 | 19.03 ± 0.30 | 20.47 ± 0.30 |
| ***Ifng*** | 31.46 ± 0.18 | 32.95 ± 0.33 | 31.15 ± 0.37 |
| ***Il10*** | 33.08 ± 0.20 | 26.52 ± 0.37 | 30.51 ± 0.46 |
| ***Gapdh*** | 13.51 ± 0.13 | 12.56 ± 0.08 | 11.61 ± 0.16 |

**Table S3: Protein levels [pg/ml] from cytokine detection assay.**

| | vehicle-only | LPS | LPS + IL10 |
|---|---|---|---|
| **TNFα** | 5.29 ± 1.73 | 4073.46 ± 455.09[§] | 380.95 ± 37.74 |
| **IL6** | 12.54 ± 2.41 | 23241.02 ± 163.27[§] | 6342.21 ± 1082.86 |
| **IL1β** | 0.05 ± 0.01[*,#] | 56.41 ± 6.73 | 4.00 ± 0.45 |
| **IFNγ** | 0.02 ± 0.01[*,#] | 0.60 ± 0.05 | 0.14 ± 0.02 |
| **IL10** | 0.79 ± 0.11[#] | 205.31 ± 14.75 | 6579.11 ± 148.24[§] |

**[*] two values with non-detectable target protein content**

**[#] values below dynamic detection range**

**[§] values above dynamic detection range**
